# GroEL-proteotyping of bacterial communities by tandem mass spectrometry

**DOI:** 10.1101/2023.07.03.546649

**Authors:** Simon Klaes, Shobhit Madan, Darja Deobald, Myriel Cooper, Lorenz Adrian

**Author notes:** corresponding author: Simon Klaes.

## Abstract

Profiling bacterial populations in mixed communities is a common task in microbiology. 16*S* rRNA-based sequencing is a widely accepted and functional approach but relies on amplification primers and cannot quantify isotope incorporation. Tandem mass spectrometry proteotyping is an effective alternative for taxonomically profiling microorganisms. We suggest that targeted proteotyping approaches can complement traditional population analyses. Therefore, we describe an approach to assess bacterial community compositions at the family level using the taxonomic marker protein GroEL, which is ubiquitously found in bacteria, except few obligate intracellular species. We refer to our method as GroEL-proteotyping. GroEL-proteotyping is based on high-resolution tandem mass spectrometry of GroEL peptides and identification of GroEL-derived taxa via a Galaxy workflow and a subsequent Python-based analysis script. Its advantage is that it can be performed with a curated and extendable sample-independent database and that GroEL can be pre-separated by SDS-PAGE to reduce sample complexity, improving GroEL identification while simultaneously decreasing the instrument time. GroEL-proteotyping was validated by employing it on a comprehensive raw data set obtained through a metaproteome approach from synthetic microbial communities as well as real human gut samples. Our data show that GroEL-proteotyping enables fast and straightforward profiling of highly abundant taxa in bacterial communities at reasonable taxonomic resolution.

## 1 Introduction

Bacterial communities govern our planet, impacting essential ecosystem services such as soil fertility [1], carbon dioxide fixation [2], bioremediation [3], and wastewater treatment [4]. One aspect of understanding the functionality of microbial communities is the identification of their microbial composition. Commonly, microbial compositions are described by (high-throughput) sequencing of 16*S* small subunit ribosomal-RNA (16*S* rRNA) gene amplicons. Several 16*S* rRNA-gene-based fingerprinting techniques, such as denaturing gradient gel electrophoresis (DGGE) [5] and terminal restriction fragment length polymorphism (T-RFLP) analyses [6], have been developed to allow the monitoring of the microbial community dynamics at reasonable costs and speed. However, all these 16*S* rRNA gene-based methods are PCR-dependent and introduce a primer bias [7]. Additionally, the variability and multiplicity of the 16*S* rRNA gene as well as the low taxonomic resolution of its short-reads, can distort bacterial community analyses [8,9]. Analyzing several hypervariable regions of the 16*S* rRNA gene [10] or other taxonomic marker genes, such as *rpoB* [11] or *groEL* [12], can enhance taxonomic resolution.

Metaproteomics is the mass spectrometric analysis of proteins from microbial communities [13]. Given the central role of proteins in metabolic processes, metaproteomics serves as a tool to investigate the functional dynamics of microbiomes, shedding light on microbial networks and their interplay with the environment [14–16]. In combination with isotope labeling, metaproteomics can link microbial populations to physiological activities, *e.g.*, to the use of a specific carbon source [17], and monitor uptake, incorporation, and interspecies transfer of isotopically labeled substrates by protein-based stable isotope probing (protein-SIP) [18]. Metaproteomic datasets are typically extensive and can be used to derive species-specific taxonomic profiles (proteotyping) within microbiomes based on all identified proteins [19–21]. In the classical proteotyping approach, a sample-specific protein database derived from (meta)genome sequencing is used for the accurate identification and quantification of proteins from mass spectrometric raw data [22,23]. When metagenomics data is not available, alternatives like RNA sequencing data [24] or universal, broad-spectrum databases such as NCBInr or UniProtKB/Swiss-Prot can be employed [25–30]. However, the large size of broad-spectrum databases leads to extended computation times, potentially hampering protein identification via target-decoy strategies [31]. Iterative cascade searches can reduce the effect of hampered protein identifications but may at the cost of increased computation times [32,33].

As an alternative, initial taxonomic profiling can assist in constructing comprehensive sample-specific databases for subsequent functional analyses [34]. Lower-resolution taxonomic profiles can be obtained using universal reference databases of taxonomic marker proteins and focusing only on a small subset of a comprehensive metaproteomic dataset. Specifically, highly abundant proteins, such as GroEL, translation initiation factor 2, elongation factors, or ribosomal proteins allowed profiling microbial communities at order level in human gut samples [34], and at domain level in mock assemblies and soil samples [35]. Utilizing a universal reference database of taxonomic marker proteins offers the advantage of eliminating the need to generate sample-specific metagenomic databases. This labor-intensive process requires repetition for each new sample site, albeit at the cost of potentially limiting the depth of functional insights. Furthermore, taxonomic marker databases tend to be smaller compared to universal, broad-spectrum databases, resulting in reduced computation time requirements.

Our research aimed to develop and evaluate a workflow enabling a semiquantitative characterization of microbial communities with high taxonomic resolution utilizing tandem mass spectrometric data focusing on GroEL as a taxonomic marker protein for bacteria. To achieve this objective, we created a sample-independent GroEL database for peptide identification and developed a Python script that facilitates protein and taxonomy inference. Our approach was evaluated by applying it to synthetic microbial communities and metaproteome data obtained from the human gut, ensuring its robustness and applicability across different microbial ecosystems.

## 2 Materials and methods

### 2.1 Cultivation, harvesting, and protein extraction

All cultivations were done in 100 mL cotton plugged Erlenmeyer flasks with 50 mL of nitrate-free DSMZ 586 medium at 30 °C in the dark on a rotary shaker under aerobic conditions. *Thauera aromatica* K172 and *Pseudomonas putida* KT2440 were separately grown and harvested during the early stationary phase by centrifuging at 16 000 x g for 10 min. Cell pellets were washed with 50 mM ammonium bicarbonate (AMBIC) buffer, pH 7.9. For the co-cultivation of *P. putida* and *T. aromatica*, we first inoculated the medium with approximately 10^6^ cells mL^-1^ of *T. aromatica*. After 15 h of incubation, we injected approximately 10^5^ cells mL^-1^ of *P. putida*. Samples were taken directly before adding *P. putida* and after 0; 18; 25; 39; and 48 h incubation time. Cells were harvested from 1 mL samples by centrifugation at 16 000 x g for 10 min, and the pellet was washed with 50 mM AMBIC buffer. Cell pellets were stored at -80 °C until further analysis. For protein extraction, cell pellets were resuspended in 50 mM AMBIC buffer and lysed by three freeze and thaw cycles, utilizing liquid nitrogen and a thermal shaker operating at 40 °C. Additionally, a 30 s treatment in a sonication bath was applied to enhance lysis efficiency. Cell debris and insoluble proteins were removed by centrifuging at 16 000 x g for 10 min. Protein concentrations were determined with the enhanced protocol of the bicinchoninic acid (BCA) assay kit (Pierce, Thermo Scientific). For synthetic bicultures, crude extracts of *T. aromatica* and *P. putida* were mixed in varying protein ratios, resulting in total protein content of 5 µg in 30 µL AMBIC buffer.

### 2.2 Sample preparation for protein mass spectrometry

Crude extract samples containing 5 µg protein in 30 µL AMBIC buffer were first amended with 40 ng bovine serum albumin (BSA) as a quality control measure. Then, the samples were sequentially treated with 62.5 mM dithiothreitol and 128 mM 2-iodoacetamide to reduce and alkylate cysteine residues. Subsequently, proteins were digested overnight with 0.63 µg trypsin (Promega). Tryptic digestion was stopped by adding formic acid to a final concentration of 1.8% (v/v). Undigested and precipitated proteins were removed by centrifugation at 16 000 x g for 10 min, and samples were dried by vacuum centrifugation. Subsequently, the peptides were resuspended in 0.1% (v/v) formic acid and desalted using C_18_ Zip tips (Pierce, Thermo Scientific). The peptides were again dried by vacuum centrifugation before being reconstituted in 50 µL of 0.1% (v/v) formic acid for nano-liquid chromatography coupled to tandem mass spectrometry (nLC-MS/MS) analysis.

In separate experiments, we decreased the heterogeneity of the crude extracts before nLC-MS/MS analysis by sodium dodecyl sulfate-polyacrylamide gel electrophoresis (SDS-PAGE). For SDS-PAGE, the crude extract containing 5 µg protein in 30 µL AMBIC buffer was mixed with 10 µL SDS reducing buffer and incubated at 95 °C for 10 min to fully denature proteins. After cooling to room temperature, samples were subjected to SDS-PAGE at 110 V for 60–90 min and Coomassie stained. Protein bands with a molecular weight of approximately 60 kDa, corresponding to the size of GroEL, were excised. The gel slices were destained with acetonitrile. Then, proteins captured within the gel slice were reduced and alkylated by sequential incubation in 50 µL of 10 mM dithiothreitol and 50 µL of 100 mM 2-iodoacetamide for 60 min each. Subsequently, 40 ng of reduced and alkylated BSA was added as a quality control measure. Proteins were then digested with 0.1 μg trypsin (Promega) at 37°C overnight. After digestion, peptides were extracted from gel pieces with 50% (v/v) acetonitrile and 5% (v/v) formic acid and dried by vacuum centrifugation. Next, peptides were resuspended in 20 µL of 0.1% (v/v) formic acid, desalted using C_18_ Zip tips (Pierce, Thermo Scientific), and repeatedly dried by vacuum centrifugation. Finally, the desalted peptides were resuspended in 50 µL of 0.1% (v/v) formic acid for nLC-MS/MS analysis.

### 2.3 Mass spectrometry

Desalted peptides were separated on an UltiMate 3000 RSLCnano high-performance nano-UPLC system (Thermo Scientific) coupled to an Orbitrap Fusion Tribrid mass spectrometer (Thermo Scientific) via a TriVersa NanoMate nano-electrospray ionization (nano-ESI) ion source (Advion). For in-solution digested samples, 3 µL of the peptide solution was injected, whereas 5 µL was injected for in-gel digested samples. Peptides were first loaded for 3 min onto the Acclaim PepMap 100 C_18_ trap column (75 µm x 2 cm, 3 µm material, Thermo Scientific) with 5 µL min^-1^ of 3.2% (v/v) acetonitrile in water at 0.1% formic acid. Then, the trap column was switched into line with the Acclaim PepMap 100 C_18_ analytical column (75 µm x 25 cm, 3 µm material, Thermo Scientific) heated up to 35 °C to separate peptides at a flow rate of 0.3 μL min^-1^ using a gradient of 145 min (for in-solution digested samples) or 60 min (for in-gel digested samples) from 3.2% to 72% (v/v) acetonitrile in water at 0.1% (v/v) formic acid. The ion source operated in positive mode at a spray voltage of 2.4 kV and a source temperature of 275 °C. The mass spectrometer run in data-dependent mode with a cycle time of 3 s. Internal mass calibration was performed using a lock mass of 445.12003 *m/z*. Precursor ions were scanned in the Orbitrap mass analyzer over a range of 350–2000 *m/z* with a resolution of 120 000, automatic gain control (AGC) target of 4 x 10^5^ ions, and a maximum injection time of 50 ms. Precursor ions with a minimum intensity of 5 x 10^4^ and charge state of +2 and +3 (for in-solution digested samples) or +2 to +7 (for in-gel digested samples) were selected and isolated by the quadrupole in a window of 1.6 *m/z* accumulating to an AGC target of 5 x 10^4^ ions with a maximum injection time of 54 ms (for in-solution digested samples) and 120 ms (for in-gel digested samples). The isolated precursor ions were fragmented using higher energy collisional dissociation (HCD) at 30% relative collision energy. Fragment ions were scanned with the Orbitrap mass analyzer at a resolution of 30 000 (for in-solution digested samples) or 60 000 (for in-gel digested samples), respectively. Precursor ions with the same mass (±10 ppm) were excluded for further precursor selection for 30 s.=

### 2.4 Databases for mass spectrometric analysis

We assembled a comprehensive dataset of bacterial GroEL protein sequences by downloading all available amino acid sequences from the National Center for Biotechnology Information (NCBI) in GenBank format on September 7, 2021 [36]. The dataset was used to generate a file in fasta format as input for peptide identification search engines (see below) and to generate a second file with tab-separated values (tsv) for protein and taxonomy inferences. The tsv-database included in separated columns accession number, amino acid sequence, expected tryptic peptides, and taxonomic classification at various levels (kingdom, phylum, class, order, family, and genus). Tryptic peptides were calculated with pyOpenMS [37], accepting two missed cleavages and a peptide length of 6–144 amino acids. The taxonomic classification was updated by cross-referencing the “List of Prokaryotic names with Standing in Nomenclature” (LPSN) on October 18, 2022 [38]. We also included GroEL sequences from the separately published genomes of *Roseobacter* sp. AK199 and *Chromobacterium violaceum* CV026 [20]. The common Repository of Adventitious Proteins (cRAP) was appended to the database to identify common contaminants. Additionally, we downloaded the whole proteome databases of *T. aromatica* K172 (CP028339.1) and *P. putida* KT2440 (NC_002947.4) from NCBI. The detectability of GroEL peptides by tandem mass spectrometry was assessed using DeepMSPeptide [39].

### 2.5 Analysis of mass-spectrometric raw data files

Thermo raw files were first converted into mzML format using msConvert (ProteoWizard) [40]. Next, we analyzed the mzML files using a customized MetaProSIP workflow [41] on the Galaxy platform [42]. Briefly, peptides were identified using MS-GF+ [43] and MetaProSIP [44]. We accepted two missed trypsin cleavages, peptide lengths of 6–40 amino acids, and precursor *m/z* deviations of 5 ppm. Oxidation of methionine and carbamidomethylation of cysteine were set as dynamic and static modifications, respectively. The false discovery rate of identified peptide sequences was kept below 0.01, based on the SpecEValue calculated by MS-GF+. The peptide-centric output of MetaProSIP was used as input for a custom GroEL-proteotyping Python script. The isobaric amino acids leucine (L) and isoleucine (I) were treated as indistinguishable. Peptides listed in the cRAP database were excluded from the analysis with the customized GroEL-proteotyping Python script. The Python script assigns detected peptides to each possible GroEL protein in the tsv-database based on matching sequences. GroEL proteins with the same set of detected peptides are merged into protein groups and sorted by the number of detected peptides in descending order. For each GroEL protein group, only the detected peptides not present in another GroEL protein group containing more peptides are counted, resulting in a Top Rank Count. GroEL protein groups are then filtered by the Top Rank Count with a threshold (≥ 5), and the taxonomy of the remaining GroEL protein groups is read from the tsv-file. For each taxonomic level, the most frequent taxonomic description and its frequency within the GroEL protein group are calculated. The most frequent taxonomic description is used to merge GroEL protein groups into taxonomic groups at the level of interest, which are then quantified by the sum of the MS1 precursor ion intensities (based on peak area) or the number of non-redundant peptides within the taxonomic group. In the current version of the Python script (version 1.0.0), non-redundant peptides are only counted once for the same taxonomic group and multiple times for different taxonomic groups. Furthermore, the precursor intensity of peptides assigned to multiple taxonomic groups is distributed proportionately to the total number of peptides assigned to each group. To evaluate our workflow, we used published metaproteome data from synthetic microbial communities (PRIDE repository PXD006118). Specifically, we used the raw data acquired with an LC gradient of 260 min (run 4 and 5) as this provided sufficient data for community analysis [20]. The identified peptides of technical replicates were pooled before running our GroEL-proteotyping Python script. Furthermore, we applied our analysis workflow to a metaproteome dataset of human gut microbiota (PRIDE repository PXD020786) [45]. Identified peptides of biological replicates were pooled before running the GroEL-proteotyping Python script to allow comparison with the original study (Supplementary Table 3 of García-Durán et al. [45]).

### 2.6 Data availability

The mass spectrometric raw data and protein sequence databases have been deposited to the ProteomeXchange Consortium (http://proteomecentral.proteomexchange.org) via the PRIDE partner repository [46] with the data set identifier PXD043106 (Username; Password: KzeQ5W8r). The GroEL-proteotyping Python script is publicly available on GitLab (https://git.ufz.de/klaes/groel-proteotyping/). Any other relevant data will be made available upon request.

## 3 Results

### 3.1 Establishing a GroEL database and analyzing protein and peptide sequences

Our download of bacterial GroEL sequences from NCBI in September 2021 resulted in 284 351 GroEL homologous sequences, of which 72 759 were non-redundant. We restricted the download to sequences of a length of 6–1500 amino acids, resulting in a median length of 542 amino acids in the database. Prediction of trypsin digestion sites in the non-redundant protein sequences allowing up to two missed cleavages, a length of 6–144 amino acids, and no ambiguous amino acid codes (B, J, O, U, X, or Z) resulted in 9 091 897 peptides, of which 1 875 469 were non-redundant.

To evaluate the taxonomic relevance of GroEL-derived peptides, we calculated their taxon-specificity across different taxonomic levels. Additionally, we predicted their MS-detectability using DeepMSPeptide [39]. For this purpose, we initially standardized taxon names in our database according to the nomenclature provided by the “List of Prokaryotic names with Standing in Nomenclature” (LPSN). Taxon names matching the LPSN nomenclature were kept, while taxon names not matching the LPSN nomenclature were excluded. Subsequently, peptides exclusively attributed to a single taxon were counted as taxon-specific peptides, with leucine (L) and isoleucine (I) treated as indistinguishable. Our results show a decline in the count of peptides with standardized taxon names, the number of taxon-specific peptides, and their predicted detectability with increasing taxonomic resolution (Table 1).

**Table 1.** Absolute and relative numbers of peptides with a standardized taxon name according to LPSN and taxon-specific peptides in the GroEL database at different taxonomic levels.

| Number of non-redundant GroEL peptides | Taxonomic level |  |  |  |  |
| --- | --- | --- | --- | --- | --- |
|  | Phylum | Class | Order | Family | Genus |
| Standardized taxon name<sup>a</sup> | 1 798 737<br>(95.9%) | 1 564 290<br>(83.4%) | 1 410 836<br>(75.2%) | 1 343 631<br>(71.6%) | 1 315 614<br>(70.1%) |
| Taxon-specific<sup>b</sup> | 1 732 937<br>(96.3%) | 1 485 993<br>(95.0%) | 1 233 553<br>(87.4%) | 1 205 161<br>(89.7%) | 1 122 799<br>(85.3%) |
| Detectable by tandem mass spectrometry<sup>c</sup> | 690 492<br>(39.8%) | 587 287<br>(39.5%) | 484 600<br>(39.3%) | 470 112<br>(39.0%) | 435 776<br>(38.8%) |

<sup>a</sup>: Relative numbers based on the total number of non-redundant peptide sequences.
<sup>b</sup>: Relative numbers based on the number of peptides affiliated with an organism with a standardized taxon name.
<sup>c</sup>: Relative numbers based on the number of taxon-specific peptides. Detectability was predicted using DeepMSPeptide [39]

### 3.2 Evaluating the sensitivity and specificity of GroEL-proteotyping and its protein filtering routine

To evaluate the performance of our database in detecting GroEL peptides, we analyzed pure cultures of *T. aromatica* K172 and *P. putida* KT2440. In this regard, each microorganism was separately cultivated, followed by protein extraction and tryptic digestion. The resulting peptides were analyzed by mass spectrometry and evaluated against two databases: *i)* a database representing the full proteome of *T. aromatica* K172 or *P. putida* KT2440, respectively, and *ii)* our GroEL database.

When comparing our data against the whole proteome databases, the total number of identified peptides was higher for *P. putida*, but the count of identified GroEL peptides was lower compared to *T. aromatica* (Table 2). When using the GroEL database instead of the whole proteome database, we observed a slight decrease in the number of detected GroEL peptides for both strains. However, the average precursor peak intensity for GroEL peptides was similar for both organisms. In summary, this initial experiment showed that in pure cultures, GroEL peptides of *T. aromatica* K172 exhibited stronger signal intensities than GroEL peptides of *P. putida* KT2440. Furthermore, the use of our GroEL database resulted in a slightly decreased detection of GroEL peptides.

**Table 2.**
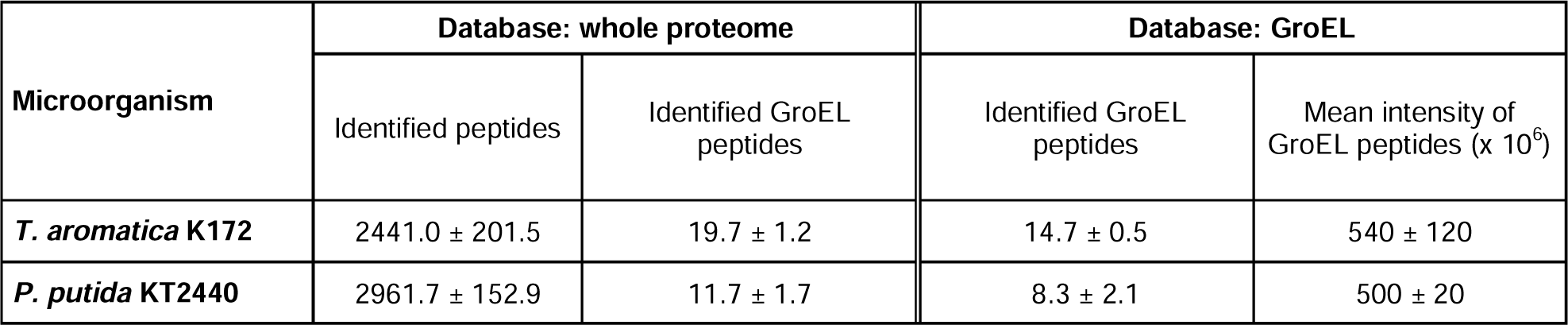
Peptides identified from protein extracts of *Thauera aromatica* K172 and *Pseudomonas putida* KT2440. Peptides were identified using the whole proteome database of *T. aromatica* K172 (“K172”) containing 3 335 entries or *P. putida* KT2440 (“KT2440”) with 5 450 entries, respectively, or the GroEL database with 72 759 non-redundant entries. Values represent means of biological triplicates ± standard deviations.

| Microorganism | Database: whole proteome |  | Database: GroEL |  |
| --- | --- | --- | --- | --- |
| | Identified peptides | Identified GroEL peptides | Identified GroEL peptides | Mean intensity of GroEL peptides ($\times 10^6$) |
| *T. aromatica* K172 | 2441.0 ± 201.5 | 19.7 ± 1.2 | 14.7 ± 0.5 | 540 ± 120 |
| *P. putida* KT2440 | 2961.7 ± 152.9 | 11.7 ± 1.7 | 8.3 ± 2.1 | 500 ± 20 |

To evaluate the applicability of GroEL peptide mass spectrometry for relative quantification of subpopulations, we experimented with a synthetic mixture of *T. aromatica* K172 and *P. putida* KT2440. Initially, each of the two strains was cultivated separately. Subsequently, proteins were extracted from them and crude extracts of *T. aromatica* K172 and *P. putida* KT2440 were mixed in predetermined protein ratios, ensuring a total protein mass of 5 µg. Before nLC-MS/MS analysis, GroEL was separated from other proteins by SDS-PAGE, decreasing the heterogeneity of the sample, which resulted in a 1.8-fold increase in the number of detected GroEL peptides. Peptides often can be assigned to multiple proteins, known as the protein inference problem. Furthermore, large databases tend to increase the number of false-positive detections. To remove false-positive detections and to infer proteins and taxonomy unambiguously, we developed a custom Python script that filters GroEL protein groups by the Top Rank Count, which only counts peptides once for the largest GroEL protein group (for the description of this filtering routine, see Materials & Methods).

Various Top Rank Count thresholds were systematically evaluated to eliminate false positive identifications while retaining the ability to detect low-abundant organisms (Figure 1). The implementation of a Top Rank Count threshold ≥ 5 eliminated all false-positive identifications at the genus level. Notably, both organisms were consistently detected at the genus level in all biological replicates at different protein ratios, demonstrating a relative detection limit of 1% of total protein content for low abundant organisms. Unfortunately, quantification based on the number of detected GroEL peptides yielded relative abundance estimations for the two genera close to 50% across all ratios, failing to reflect the actual mixing ratios. Quantification based on the sum of the precursor intensities from detected GroEL peptides provided more accurate estimations. In mixtures containing 1%, 5%, and 10% *T. aromatica K172* proteins, we observed *Thauera* abundance values of 11.9% ± 3.1%, 8.9% ± 1.1%; and 22.4% ± 0.6% respectively.

**Figure 1.**
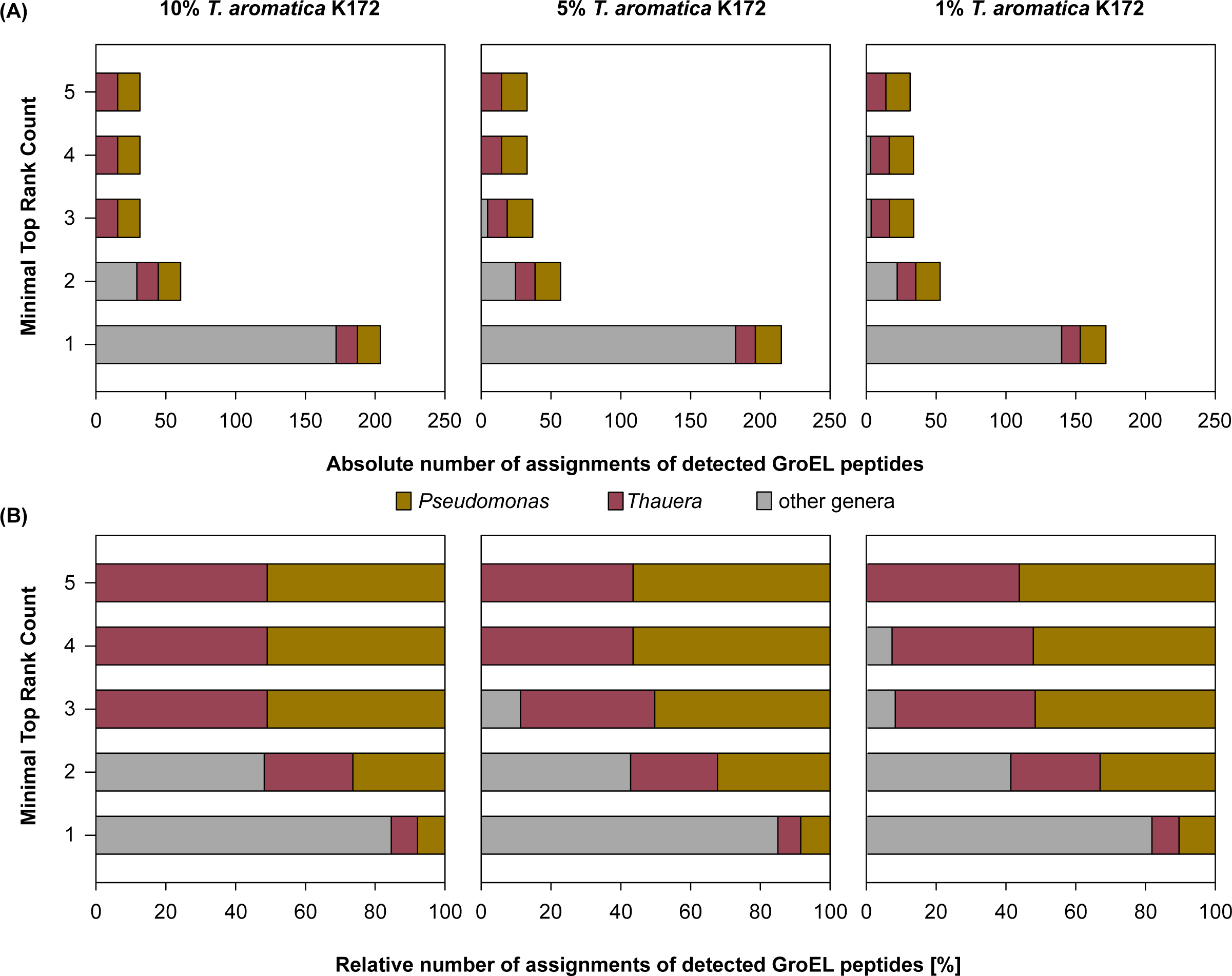
Evaluating sensitivity **(A)** and specificity **(B)** of GroEL-proteotyping in detecting organisms at the genus level in mixed protein extracts. Mixed protein extracts were composed of *Thauera aromatica* K172 and *Pseudomonas putida* KT2440 proteins in different ratios, and different filtering criteria were applied. Bars represent means of three biological replicates.

Taken together, our findings indicate that GroEL-proteotyping allows for the differentiation of two organism mixtures at the genus level, achieving a relative detection limit of 1% of the total protein content, representing approximately 50 ng protein in the crude extract. Quantification based on the sum of the precursor intensities proved to be more reliable and closer to the actual mixing ratios than GroEL peptide count-based quantification. However, in all mixtures, we consistently detected a higher relative abundance of *Thauera* than originally added.

### 3.3 GroEL-proteotyping in action I: Reanalyzing raw data of synthetic microbial communities

In the previous experiments, communities of only two bacteria were investigated. To examine if the filtering and quantification techniques are applicable for characterizing more complex bacterial consortia, we reanalyzed proteomic raw data files of three synthetic microbial communities originally assembled by Kleiner *et al.* [20]. The synthetic communities comprised crude protein extracts of one archaeon, one eukaryote, five bacteriophages, three strains of Gram-positive bacteria, and 18 (Mock A and B) or 22 (Mock C) strains of Gram-negative bacteria mixed in different ratios (Supplementary Tables 1–3 of Kleiner et al. [20]). The bacterial population contained 12 (Mock A and B) to 15 (Mock C) bacterial families. Since our method is specialized in bacterial community characterization, we focused only on these bacterial families.

We again investigated different thresholds of the Top Rank Count to eliminate false-positive identifications while still detecting low-abundance microorganisms (Figure 2). Filtering by a Top Rank Count threshold ≥ 5 (as described above) resulted in some false identifications at the genus level. However, at the family level, this filter effectively eliminated all incorrect identifications while maintaining an acceptable sensitivity for detecting low-abundance organisms. Therefore, we used this threshold for subsequent community analysis at the family level.

**Figure 2.**
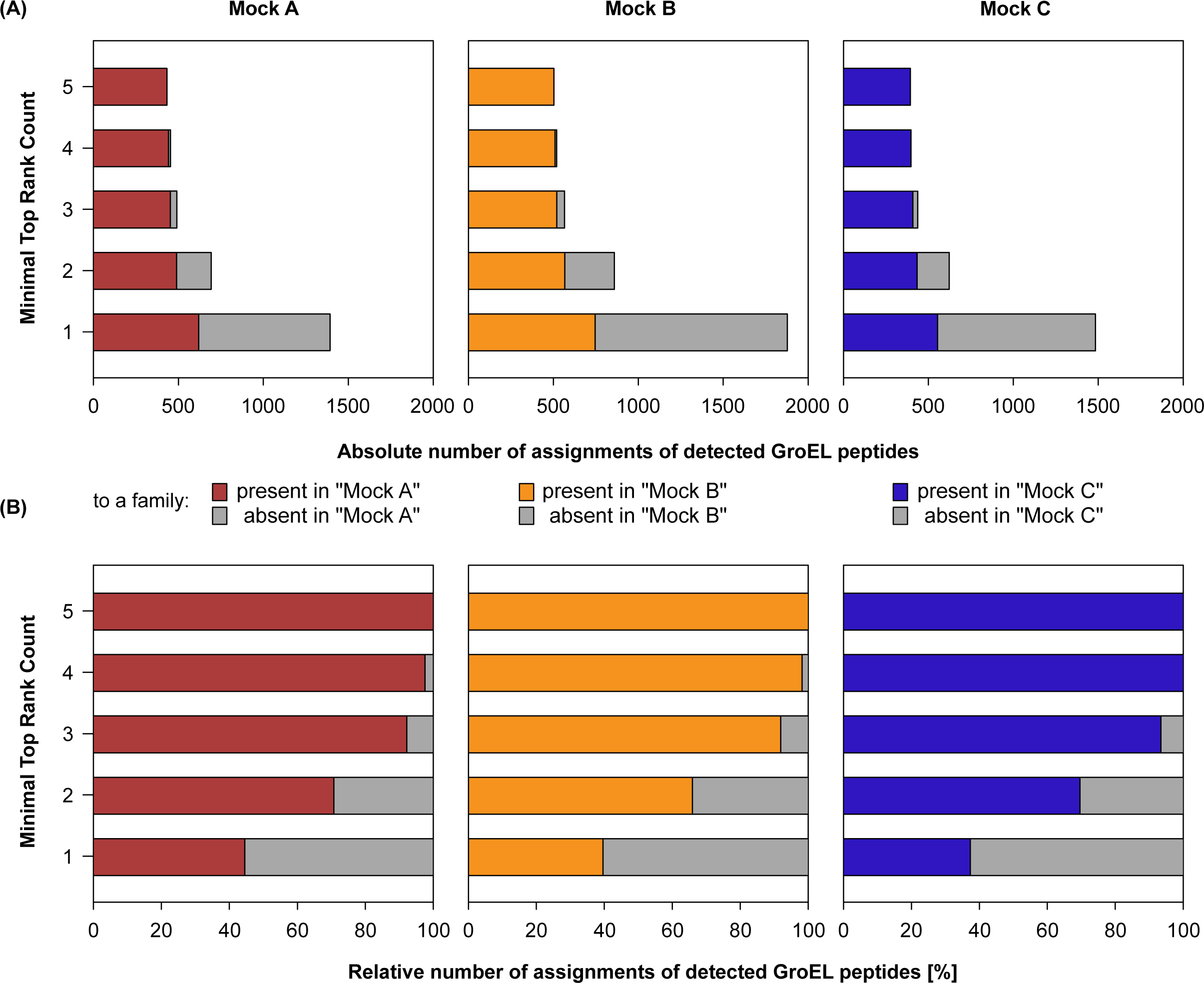
Evaluating sensitivity **(A)** and specificity **(B)** of GroEL-proteotyping using different Top Rank Count filtering thresholds in detecting peptides assigned to bacterial families that were present in three synthetic microbial communities (Mock A–C) assembled by Kleiner *et al.* [20]. The detected peptides were assigned to specific GroEL proteins. GroEL proteins with the same set of detected peptides were merged into GroEL groups, which were then filtered by the Top Rank Count ≥ 5. Bars represent the means of four biological replicates.

In the raw data files of the three synthetic communities assembled by Kleiner *et al.* [20], we detected all bacterial families that were present at a relative bacterial protein content above 2.8%, while no absent family was falsely detected (Figure 3A). However, the detection of scarce families was inconsistent between the assessments of the different synthetic communities. Specifically, we detected all bacterial families in all four biological replicates of Mock B, which has the most balanced actual protein distribution of 4.8–19.1% of the total protein content for each family (Figure 3A, yellow). In Mock A, all families except the lowest-abundant family *Staphylococcaceae*, accounting for 0.7% of the total bacterial protein content, were detected (Figure 3A, red). On the other hand, in Mock C, which had the highest number of low-abundant bacterial families (relative bacterial protein content ≤ 2.8%), and the most uneven actual protein distribution, with families representing 0.2–41.8% of the total bacterial protein content, we detected all highly abundant bacterial families (relative bacterial protein content > 2.8%) (Figure 3A, blue). However, the detection of low-abundant families with relative bacterial protein content ≤ 2.8% was inconsistent: we detected *Chromobacteriaceae* (1.3%), *Desulfovibrionaceae* (1.0%), *Rhodobacteraceae* (1.0%), *Roseobacteraceae* (1.7%), and *Thermaceae* (1.8%), but not *Staphylococcaceae* (2.8%), *Alteromondaceae (1.0%)*, *Bacillaceae (0.8%)*, *Nitrosomonadaceae (0.7%)*, and *Nitrospiraceae (0.2%)* (Figure 3A, blue).

**Figure 3.**
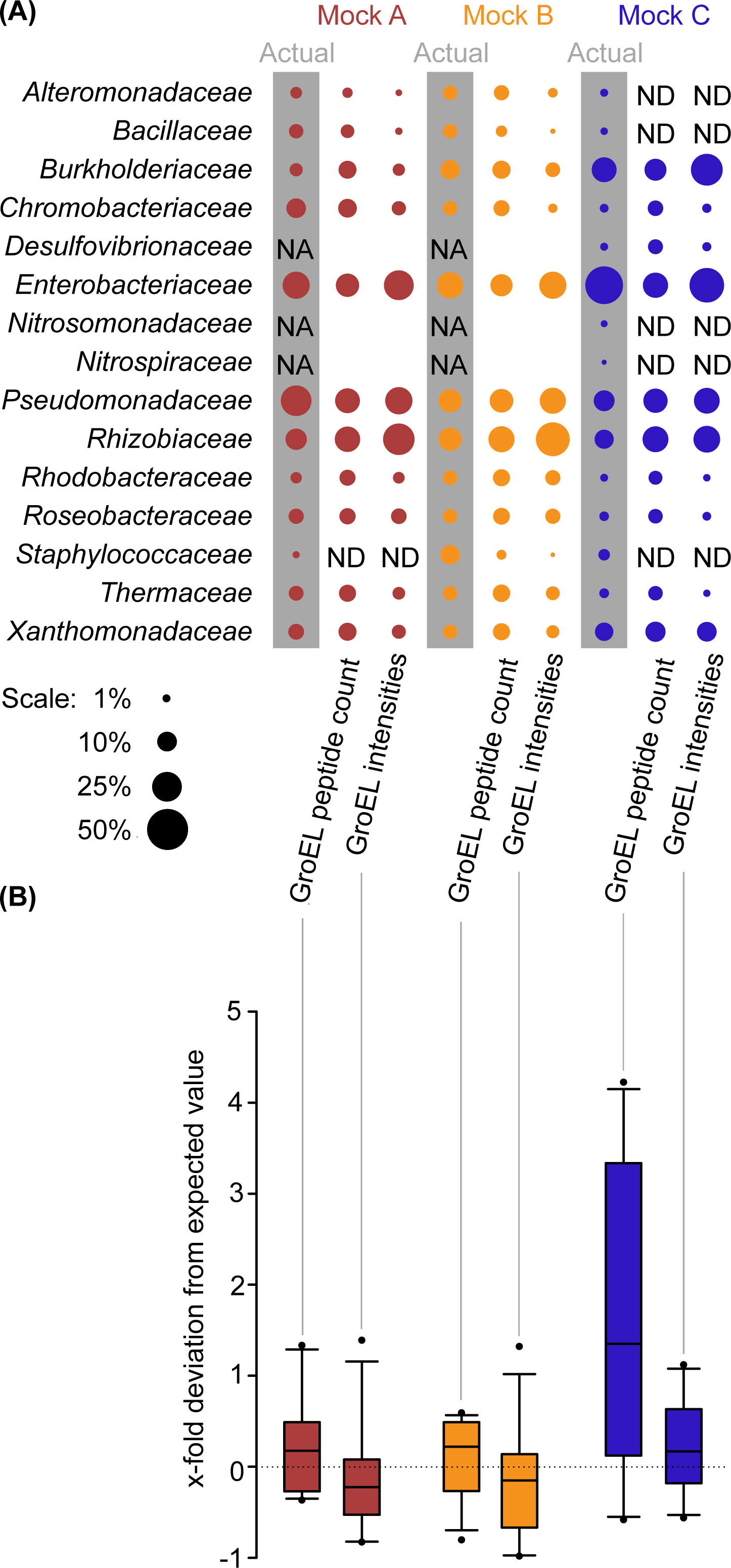
Profiling the taxonomic composition of bacterial subpopulations in synthetic microbial communities using GroEL-proteotyping. The metaproteome dataset of synthetic communities was obtained from Kleiner *et al.* [20]. We used the same graphic layout to facilitate comparison. **(A)** Comparison of actual community shares (shaded values) with GroEL-based quantification: ‘Peptide number’ – quantification based on the number of non-redundant GroEL peptides assigned to that family; ‘Intensities’ – quantification based on the sum of the precursor intensities of the detected GroEL peptides. Quantities are represented as bubble areas. The figure includes information on families that were not added (NA) or not detected (ND) with our method. Results are means of four replicates. **(B)** Comparison of two quantification methods for GroEL peptides to the actual protein abundance of each family. 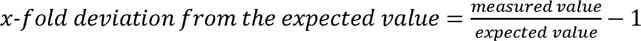. Boxes show the 1^st^ and 3^rd^ quartile, the median, and the whiskers indicating the 10^th^ and 90^th^ percentile, with filled circles representing outliers. ND were removed before plotting. A value of 0 is depicted as a dotted line, indicating equal measurement and input. Negative values indicate underestimation, while positive values indicate overestimation.

In addition, we compared two quantification methods: *i)* analyzing the number of detected GroEL peptides and *ii)* analyzing the sum of the precursor intensities of detected GroEL peptides (Figure 3B). Both methods performed similarly for Mock A and Mock B, with the medians centering around zero, indicating high agreement of our analysis and input (Figure 3B, red and yellow). However, differences between the two quantification methods became apparent for Mock C. The method based on peptide counts overestimated most families (*e.g.*, *Thermaceae*) while underestimating others (*e.g.*, *Enterobacteriaceae*) (Figure 3B, blue). The deviation of the measured abundance from the actual input was smaller for the method based on precursor intensities than the method based on peptide counts. Overall, we show that the characterization of complex synthetic communities by GroEL-proteotyping is robust at the family level and more consistent when based on the sum precursor intensities of detected GroEL peptides than when based on peptide counts.

### 3.4 GroEL-proteotyping in action II: Reanalyzing raw data of human gut microbiomes

To test the applicability of GroEL-proteotyping for characterizing complex bacterial communities, we reanalyzed a gut metaproteomic dataset previously published by García-Durán *et al.* [45] pertained to six healthy individuals. We focused on the human gut microbiome because it harbors a diverse bacterial community of up to 1 150 species [47]. The original study used a protein database based on the human gut microbial gene catalog (9 878 647 sequences) [48], and human proteins from UniProt (74 451 sequences) to detect an average of 11 712 peptides per sample, 56% of which were assigned at the family level. García-Durán *et al.* identified 33 different bacterial families, of which 11 showed a relative family abundance of at least 1% (Supplementary Table 3 of García-Durán *et al.* [45]).

In our reanalysis, we used our GroEL database to detect peptides and assign them to GroEL protein groups. We filtered GroEL protein groups by the Top Rank Count with a threshold ≥ 5 and evaluated the taxonomy at the family level to achieve high specificity. Our reanalysis closely mirrored the identification of abundant families reported by García-Durán *et al.* (Figure 4) [45]. We detected all 11 families with a relative family abundance of at least 1% (in the original publication). Among the 22 families exhibiting a relative family abundance below 1%, our analysis successfully identified three families: *Streptococcaceae*, *Enterobacteriaceae*, and *Coprobacillaceae* (identified as *Erysipelotrichaceae* by [45]) (Table S1). We did not detect the remaining 19 families, identified with a relative abundance below 1% in the original publication. We observed only minor differences in the abundance of highly abundant families (at least 1% in the original publication), depending on the quantification parameter used (peptide count, peptide spectrum matches (PSMs), or sum precursor intensities). However, for a few families, we noted a substantial discrepancy between our quantification based on GroEL and the quantification based on the human gut microbial gene catalog applied in the original publication. Specifically, GroEL-proteotyping led to an overestimation of *Lachnospiraceae* and *Clostridiaceae* and an underestimation of *Bacteriodaceae*, *Eubacteriaceae*, and *Prevotellaceae* comparing to the original analysis (Figure 4).

**Figure 4.**
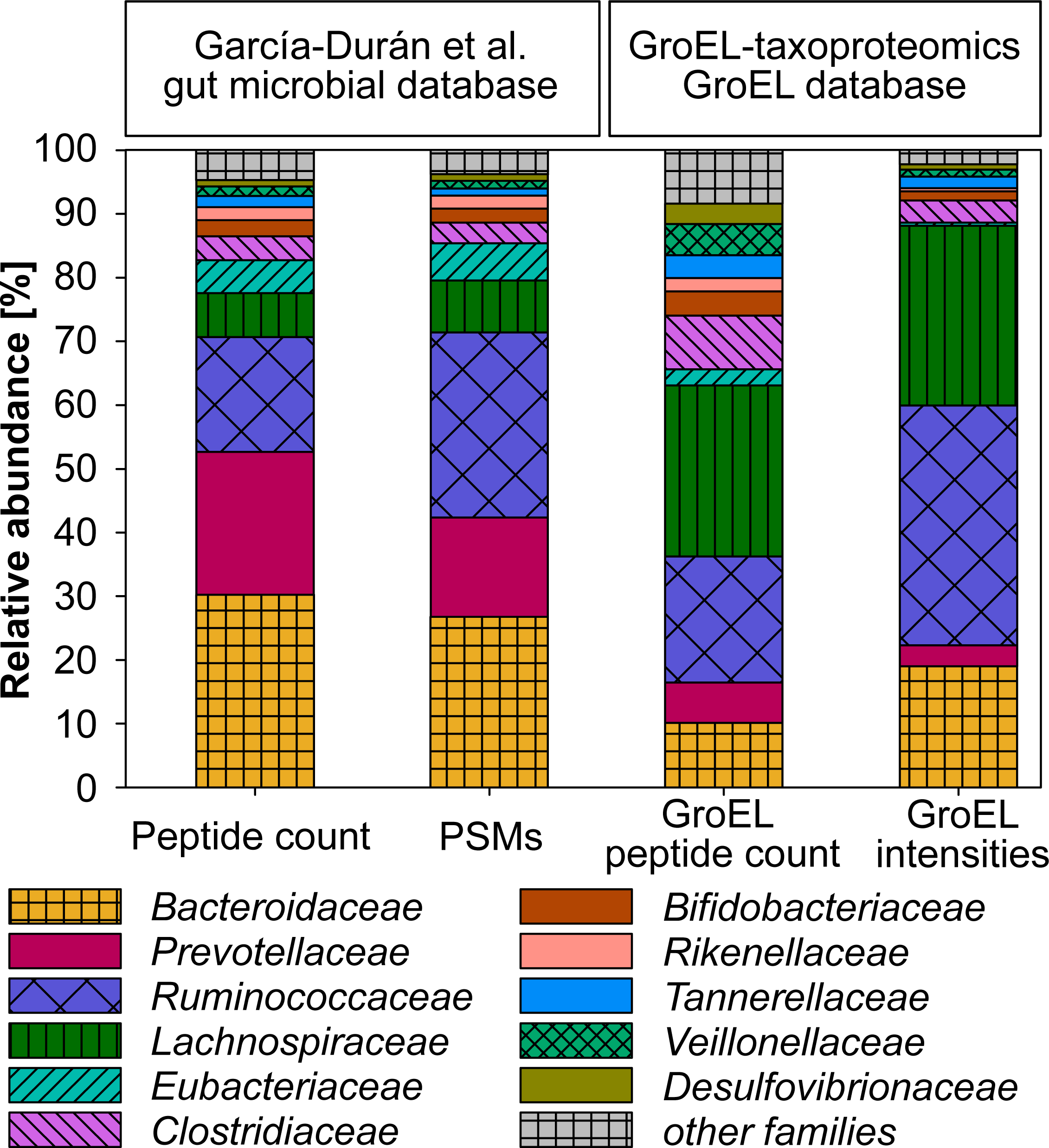
The average composition of bacterial communities at the family level, as described from the gut metaproteomic dataset of six healthy individuals by García-Durán *et al.* [45], compared with reanalysis of the same data set using GroEL-proteotyping. Peptides that could not be assigned at the family level were excluded. In the original analysis, the relative abundance of a family was determined by the number of peptide spectrum matches (PSMs) or unique peptides assigned to that family as a proportion of the total number of PSMs or unique peptides assigned at the family level, respectively. In our analysis, the relative abundance of a family was calculated based on the sum of the precursor intensities or the number of non-redundant GroEL peptides assigned to that family, relative to the total sum of the precursor intensities or the number of all filtered GroEL peptides. Families with an abundance below 1% in the original analysis were merged and represented as ‘other families’.

In the second case study, we demonstrate that GroEL-proteotyping can be applied to real and complex samples, as it yields results comparable to traditional metaproteomic approaches in detecting highly abundant bacteria. However, we also identify certain limitations associated with our method. Specifically, there are challenges in accurately quantifying and reliably detecting low-abundance bacterial families.

## 4 Discussion

Our findings show that compositions of bacterial communities can be analyzed by using GroEL as a taxonomic marker protein and a sample-independent database. This aspect holds substantial advantages, particularly in situations where rapid or continuous monitoring of community compositions is required. In our work, highly abundant families were reliably identified in all tested samples. We propose a flexible workflow (Figure 5) that can be adapted to a variety of sample preparations, nLC-MS/MS setups, peptide search engines, and quantification strategies. This workflow facilitates the analysis and can be automatized, *e.g.*, as a Galaxy workflow. It provides quantitative identifications with both, sample-independent or sample-specific databases. While the sample-independent GroEL database gave satisfactory results on the family level, a sample-specific GroEL database can improve identifications, as shown with our defined bicultures. We propose introducing the general concept of ‘targeted proteotyping’ as a distinct subcategory of proteotyping, that can be applied to different taxonomic marker proteins. Adopting the term ‘GroEL-proteptyping would then differentiate this particular method from other (targeted) proteotyping approaches.

**Figure 5.**
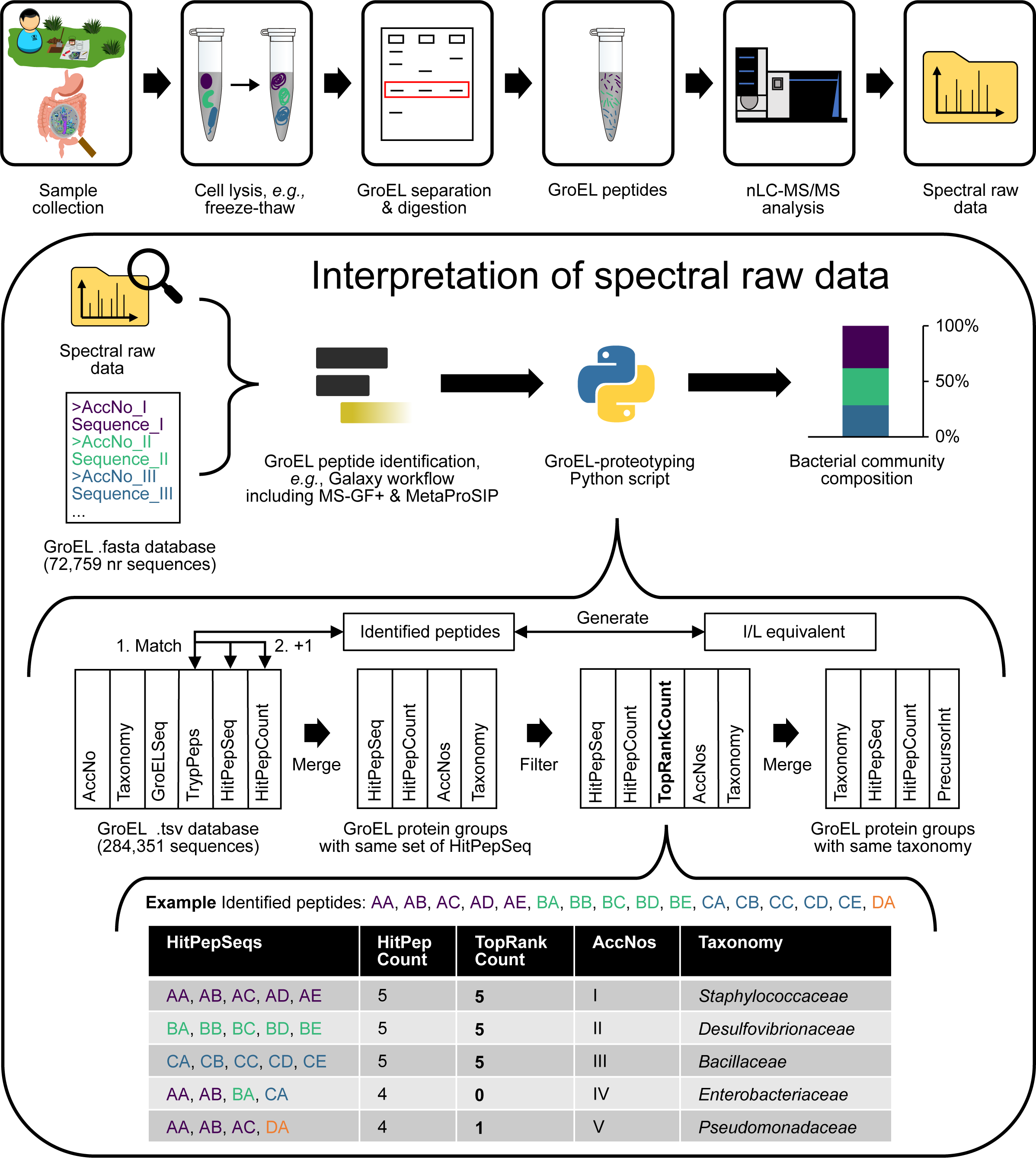
Proposed workflow for analyzing mixed bacterial communities using GroEL-proteotyping. The workflow is adaptable to various sample preparations, nano-liquid chromatography-tandem mass spectrometry (nLC-MS/MS) configurations, peptide search engines, and quantification strategies. While SDS-PAGE pre-separation of GroEL proteins can enhance the detection limit, it is not compulsory. The lower part explains the Top Rank Count approach used to filter hits. See the text for further explanations.

The analysis of our database shows that most tryptic GroEL peptides are highly taxon-specific similar to the nucleotide sequence of the *groEL* gene that has already been established as a barcode for bacteria [12]. Likewise, our workflow identified bacteria at the genus level in low-complexity samples, such as bicultures composed of *T. aromatica* K172 and *P. putida* KT2440. In more complex samples, the identification was reliable and robust at the family level. The classical metaproteomic approach allows for characterizing the same sample down to the species level [20]. However, this enhanced resolution is achieved at the expense of necessitating a sample-specific metagenomic database. In contrast, phylopeptidomics, a peptide-centric approach, achieves species-level characterization of the same sample but using a large, sample-independent NCBInr database, resulting in high computation demands [26]. Previous analyses based on GroEL or other taxonomic marker proteins without additional filtering procedures could differentiate the mock communities only at kingdom level [35]. This demonstrates the importance of effective peptide filtering and protein/taxonomy inference. Here, we achieved a higher taxonomic resolution by employing a filtering strategy for GroEL protein groups based on the number of peptides counted for top-scored proteins, which we refer to as ‘Top Rank Count’ (≥ 5). While this approach is not entirely novel, as similar filtering techniques have been implemented by Proteome Discoverer (Thermo Scientific) and MassSieve [49], its integration into our GroEL-proteotyping workflow enabled us to attain superior taxonomic resolution compared to previous GroEL-based approaches. Although GroEL-proteotyping currently has a lower taxonomic resolution and provides limited information beyond taxonomic composition, it stands out for its advantages in terms of speed and cost-efficiency compared to metaproteomics and phylopeptidomics. These benefits arise from a smaller sample-independent database and the reduced sample complexity allowing shorter instrument run times.

Taxa quantification was more robust based on the sum of the precursor intensities compared to the peptide count and a good semiquantitative estimate for the actual protein input in the complex synthetic communities. Surprisingly, none of the quantification approaches accurately reflected the relative protein amount of the strains in theIn our synthetic biculture, we consistently observed a higher signal for GroEL peptides of *Thauera* compared to those of *Pseudomonas*. One reason could be indicating that the relative GroEL expression differs between microbes, growth phases, and environmental conditions and that the difference became more apparent for the biculture setup than in the complex communities [50–52].

Our data indicate that the detection of low-abundant taxa strongly depends on the sample complexity and protein distribution across taxa. In a complex synthetic community with an uneven protein distribution, we only reliably detected bacterial families with a relative protein mass of more than 2.8%. However, pre-separation of GroEL by SDS-PAGE increased the identification of GroEL peptides 1.8-fold, resulting in a relative detection limit of 1% in low-complex bicultures. Thus, we hypothesize that using a separable marker protein allows for reducing sample complexity without losing taxonomic information, consequently enhancing the detection of low-abundant taxa. Larger protein input, pooling of gel bands, and longer LC gradients during mass spectrometry may further improve the detection of low-abundant taxa. A fast and sensitive screening of taxa present in a sample based on marker proteins could also aid in creating sample-specific databases for subsequent functional analysis of the whole metaproteomic data as shown before [34].

In our approach, the detection of organisms depends on the presence of its GroEL sequence in the database. For example, *Roseobacteraceae* was only detected in complex synthetic communities after adding the GroEL sequence of *Roseobacter* sp. AK199 to the database, demonstrating that incomplete databases bias the identification and quantification of taxa. Thus, applying GroEL-proteotyping to environmental samples containing many non-sequenced organisms is still challenging [53]. However, we successfully applied GroEL-proteotyping to human gut proteome data. Furthermore, we are confident that the rapid growth of sequence databases will massively increase database coverage. In addition, universal primers can amplify a 549–567 bp region of the *groEL* gene, allowing a targeted, fast, and sample-specific extension of the database [54].

Our study introduces targeted proteotyping as a concept for proteotyping microbial communities using taxonomic marker proteins. At present, our targeted proteotyping approach is limited to bacteria because GroEL is highly abundant in bacteria [35,55–59] (except very few intracellular *Mycoplasma* and *Ureaplasma* strains [60]), while it is only found in some archaea that most likely acquired it through horizontal gene transfer [61]. Consequently, the current scope of our approach does not encompass the detection of eukaryotes and archaea. To expand the applicability of our method to also detect eukaryotes and archaea, the incorporation of additional putative marker proteins such as ribosomal proteins, chaperonin TCP-1, or thermosome proteins should be considered.

In summary, we introduce GroEL-proteotyping as a rapid and cost-effective method for protein-based profiling of bacterial communities at the family level. In comparison to classical protein-based approaches, GroEL-proteotyping bypasses the need for sample-specific databases, saving time and reducing costs associated with database generation while achieving higher taxonomic resolution than previous targeted proteotyping approaches [34,35]. Although the implementation of our method requires access to a protein mass spectrometer, the actual analysis process, with a 60 min LC gradient, is fast, which enables the characterization of up to 20 complex samples per day, making our approach highly efficient in particular for high-throughput analyses. Furthermore, our automatable bioinformatics workflow enables the taxonomic profiling of bacterial communities within 48 hours. This can particularly be advantageous for monitoring defined or highly enriched mixed communities over time. Moreover, as the field progresses very fast, the applicability of GroEL-proteotyping will expand with increasing GroEL protein sequence databases, the development of automated workflows, the combination with stable-isotope probing, and the optimization of mass spectrometric techniques such as ion mobility devices [62].

## 5 Acknowledgments

This project was funded by the German Research Foundation (DFG) as part of the Research Training Group “Urban Water Interfaces (UWI)” (GRK 2032/2, Project H8: “Microbial transformation of mobile halogenated aromatics in redox gradients of urban hyporheic zones”). Open access was supported by the UFZ library.

We acknowledge Benjamin Scheer for assistance with mass spectrometric measurements. Furthermore, we thank Matthias Bernt and Bryson Gibbons for rapid troubleshooting regarding the Galaxy server and MS-GF+.

## 6 Author contributions

Conceptualization, Simon Klaes, Darja Deobald, Myriel Cooper and Lorenz Adrian; Data curation, Simon Klaes; Formal analysis, Simon Klaes; Funding acquisition, Myriel Cooper and Lorenz Adrian; Investigation, Simon Klaes and Shobhit Madan; Methodology, Simon Klaes, Darja Deobald and Lorenz Adrian; Software, Simon Klaes and Lorenz Adrian; Validation, Simon Klaes and Shobhit Madan; Visualization, Simon Klaes; Writing – original draft, Simon Klaes; Writing – review & editing, Simon Klaes, Shobhit Madan, Darja Deobald, Myriel Cooper and Lorenz Adrian.

